# Dynamic Changes of G-CSF etc. Nine Cytokines in Mouse Bloodstream Infection Models of *Staphylococcus aureus* and *Klebsiella pneumoniae* and their Clinical Performances

**DOI:** 10.1101/639104

**Authors:** Shang He, Ming Yang, Xinjun Li, Chen Chen, Ma Yating, Chi Wang, Jiyong Yang, Chengbin Wang

**Affiliations:** Clinical Laboratory Center, Chinese PLA General Hospital & Medical College of Chinese PLA, Beijing, 100853, China

**Keywords:** *S.aureus*, *K.pneumoniae*, Cytokine, Bloodstream infection, Dynamic Change

## Abstract

Blood culture has been considered as the gold standard to diagnose the bacterial bloodstream infection, but its long turnaround time gravely obstructed the clinical medication by physicians. Cytokines play an important role in bacterial infection. The purpose of this study was to monitor the kinetic changes of nine cytokines in mouse infection models and to infer their diagnostic value in early infection.

**Methods:** The mouse bloodstream infection model of *Staphylococcus aureus* and the other model of *Staphylococcus aureus and Klebsiella pneumoniae* were constructed respectively, and the dynamic changes of nine cytokines were monitored within 48 hours after infected with 1/2 LD_50_ bacterial concentration. Cytokines with significant differences between the two groups and PBS control group from 0 to 6 hours after infection were selected for theoretical proof in patient sera that were clearly diagnosed as bloodstream infection. Receiver operating characteristic (ROC) curve analysis was conducted to determine the clinical differentiation of different cytokines.

**Results:** Two models of *S.aureus* and *K. pneumoniae* bloodstream infection in mice were constructed successfully. In the two mouse models, six of the nine cytokines monitored were different (*P*<0.05) in each experimental group. In the 121 patient sera samples, three cytokines, IL-6, IL-12p70 and G-CSF in the infection groups and control group had showed differences. In particular, AUC of G-CSF was 0.9051, the accuracy is better than IL-6 for diagnosing the infection. In addition, only G-CSF was significantly different between the two infection groups and in the analysis of ROC curve, AUC is equal to 0.735.

**Conclusions:** G-CSF can not only judge the bacterial infection and non-infection, but also distinguish the infection of *S.aureus* from *K. pneumoniae*.

## 1. Introduction

Bacterial Bloodstream infection (BSI) is an important cause of serious morbidity and mortality. Blood cultures are regarded as the ‘gold standard’ for the detection of BSI, but it generally takes about 3-5 days and is considered as time consuming. Although MALDI-TOF mass spectrometry has gradually been introduced into clinical laboratories to reduce the time for microbial identification, the process would still take 2-3 days. Furthermore, the sensitivity of blood culture decreases significantly when antibiotic therapy had been used to treat before blood samples are taken [1, 2], or when fastidious or slow growing pathogens need to be cultured [3]. BSI often cause sepsis, which progress rapidly over the course of hours, therefore doctors must prescribe their patients receipts on empiric broad spectrum antimicrobial agents until the pathogen species and their antimicrobial susceptibility were identified. Unfortunately, this practice produces selection pressure and likely contributes to the problem of multidrug-resistant organisms and simultaneously increases the economic burden of patients [1, 2]. Therefore, fast and reliable diagnostic tools are required to provide adequate therapy in a timely manner and to enable a de-escalation of treatment[4]. It is a requirement to seek suitable biomarkers for BSI early detection in clinical settings.

When the pathogen enters the bloodstream, it first activates the body’s innate immune response [5]. The peptides, lipopolysaccharide, and teichoic acid on the surface of the bacteria may induce the pathogen-associated molecule pattern (PAMP) immune response of immune cells. The pattern recognition receptor (PRR) on the surface of innate immune cells subsequently transmits the signal to the interior of the cells, promoting or inhibiting the release of inflammatory cytokines, especially innate immune response involved in toll-like receptors (TLR). TLR plays an important role in the initial immunologic process of the bacterial infections. Therefore, inflammatory factors are potential candidates as early diagnostic markers of bacterial infection.

Up until now, number of bacterial infection markers, such as WBC, CRP, lL-6, VEGF、Eotaxin, MCP-4, and MIP-1, apresepsin, PCT, CD64, TNF-α IL-2, IL-8, IL-10 and proADM, have been described to be the potential biomarkers for BSI detection. In infections, cytokines may play key roles in the infection of pathogens [6, 7].

In order to screen clinical useful, early diagnostic markers for the bloodstream infections, we chose microorganisms with high clinical infection rate to infect mice, including eight types of bacteria (3 gram-positive bacteria and 5 gram-negative bacteria) and one type of fungi. A total of 32 cytokines or chemokines related to the bacterial infection was chosen as observation indices. We also aimed to evaluate the diagnostic biomarkers whether could distinguish the infections with gram negative bacteria from gram positive bacteria.

In this study, mice infected with *Staphylococcus aureus* or *Klebsiella pneumoniae* were used as the mouse model to monitor the dynamic variation pattern of several infection-related cytokines after infection to find potential biomarker for early diagnosis of BSI. Based on their dynamic changes in animal models in vivo, several cytokines that were different between the three groups, experimental groups of *S.aureus* and *K.pneumoniae* and healthy control group, were selected for proof verification.

## 2 Methods And Patients

### Preparation of bacteria

The standard strains *S.aureus* ATCC25923 used in the study was donated by the People’s Liberation Army 302 Hospital Laboratory. The second standard strain *K.pneumoniae* ATCC 700603 were donated by the PLA General Hospital microbiology department. The two bacterial strains were incubated in LB medium (Oxide Microbiology Products, England) for 16~18h at 37 °C. The culture was then washed twice and resuspended in sterile PBS, respectively.

### Determination of half of the lethal dose

In order to simulate the process of bacterial infection *in vivo*, mice were selected as the model of infection. To ensure that the mice were alive under the infection condition during observation and blood collection, we chose 1/2 LD50 as the dose of bacterial infections. The clinical observation and molecular biological detection were used to determine whether the mouse was in a state of infection.

Two strains of bacteria we studied were shake cultured at 37 °C in the 3 ml LB liquid medium, respectively. After 2 reactivations, the cultures were centrifuged at 1700 g for 7 min, and resuspended with sterile PBS. After repeated for 3 times, the concentration of the bacteria was quantified by Maxwell turbidity method. The resuspended bacteria were first diluted to 3.9 Maxwell’s turbidity. Then we performed a 10-fold dilution and diluted six concentrations sequentially. The diluted bacterial suspension was injected into the mice by the tail vein, and each group had 5 mice specific-pathogen-free(SPF) male CD-1(ICR) mice, with a total of 6 groups. The injection volume was 0.1ml /10 g body weight. Mouse actions, weight and death were observed and recorded daily for 7 days. The LD_50_ was calculated by Karber method.

### Animal study

A total of 105 specific-pathogen-free(SPF) male CD-1(ICR) mice, 6~8 weeks old, were purchased from the Weitonglihua (Beijing) Experimental Animal Science and Technology Co., Ltd. [SCXK (Jing) 2016-0011], weighing between (30±3) g. They were kept in a SPF facility with the temperature of 18-25°C and humidity of 50%-70%. All mice acclimated for one week before the experiment. The experimental protocol was approved by the Animal Care and Use Committee, PLA general hospital, Beijing. All experiments were performed in accordance with relevant institutional and national guidelines and regulations.

In brief, the mice were inoculated with the 1/2 LD_50_ diluent of standard strain of *S. aureus* or *K.pneumoniae* in a final volume of 0.1 mL/10g body weight via tail veins. The time of challenge was designated as time 0 of the experiment. The orbital blood of mice was collected at the 0.5h, 1h, 3h, 6h, 12h, 24h and 48h after inoculation and placed at 4°C for 8 hours, then the serum was isolated. Five repeats were set for each processing. The levels of cytokines, IL-1β, IL-5, IL-6, IL-7, IL-12p40, IL-12p70, G-CSF, IFN-γ and TNF-α, of the serum were detected using the Luminex® xMAP™ System.

Observe and record the mouse behavior, weight and death were observed and recorded at the 0.5h, 1h, 3h, 6h, 12h, 24h and 48h after inoculation.

### PCR identification of blood cultures

Blood samples were collected for culture 0.5h, 1h, 3h, 6h, 12h, 24h and 48h after injection. Cultures were streak inoculated on the columbia blood agar plate 24 hours later, and a single colony was picked for extracting DNA of *S.aureus* and. The primers of the virulence factors of the two bacteria were used to amplify corresponding DNA. The primers for *S.aureus* are: Coa F: 5’cct caa gca act the gaa aca aca3’, Coa R: 5’tga atc ttg gtc tcg ctt cat3’. The primers for *K.pneumoniae* are fimH2 F: 5’acg tgg tgg tcc cca cc3’, fimH2 R: 5’tgc cga tga tcg act gca3’.

### The verification of the experiment in human serum

In this study, 121 samples of blood were collected from the Chinese People’s Liberation Army General Hospital. 临床样本的基本情况? All blood samples we used in this study have obtained the informed consent from participants. Duplicate blood samples were also taken before treatment with antibiotics on the same day for blood culture. Samples infected by more than one type of bacterial were removed. In order to distinguish the changes in serology in healthy control group, we included a group of physical examination population as negative control. Patient recruitment criteria used for the control group were negative in blood culture, PCT <0.5 g/L, IL-6 <5.9g/L and CRP <0.8 g/L, whereas for the infection group, the requirements of positive in blood culture, PCT> 0.5g/L or IL-6> 5.9g/L or CRP> 0.8g/L were met. Among the 121 samples, 67 were *S.aureus* infected, 23 had *K.pneumoniae* and a total of 31 samples were in the control group.

### Statistical analysis

The Kolmogorov-Smirnov or Shapiro-Wilk tests were used to assess the normality of distribution of investigated parameters when appropriate. Continuous variables were described using the median (inter-quartile range [IQR]) for non-normally distributed data or the mean (standard error [SE]) for normally distributed data. Comparisons of group differences for continuous variables were made by the Mann-Whitney U test or the Student t-test as appropriate. The significance of differences in proportions was tested by the Chi-squared test. Receiver operating characteristic (ROC) curves and areas under the curves (AUCs) were calculated to evaluate the performance of each biomarker for infection. The optimal cutoff values were set for each ROC curve through the Youden Index. The values *P*<0.05 were considered statistically significant. Statistical analysis was done using SPSS v. 22.0 (software SPSS Inc., Chicago, IL, USA). All graphs were made by GraphPad Prism 7.0 software. All of ROC curves in our study were the sensitivity/specificity over the scale range computed with SPSS 22.0. The 95% confidence intervals (CIs) were calculated for all estimations.

## 3. Results

### 3.1 LD50 of the two bacteria

The LD50 of the *S.aureus* and *K.pneumoniae* was 8.1×10^8^ CFU/mL and 1.11×10^9^ CFU/mL, respectively.

### 3.2 Clinical symptoms after infection in mice

About 1 h after the injection of bacteria, the mice in the experimental group began to be of piloerection, closed eyes and decreased activity. 3 h after injection, these behaviors were more obvious, the body curled up, the mice ate less and had defecated loose stool. 24 h after injection, the mice had significantly decreased feeding and activity, eyes closed, eyes secreted more, and stools became dry. 48 h after injection, some of the mice dead, the left got back to normal condition.

### 3.3 Changes of mice weight 24 h

After injection, the mice weight of *S. aureus* group decreased by an average of 3.9 g, and the weight of *K.pneumoniae* group decreased by an average of 4.7 g. The weight of both groups maintained at low levels within 48 h and then gradually recovered. The weight of *K.pneumoniae* group recovered faster than that of the *S.aureus* group as was shown in Figure 1.

**Fig 1.**
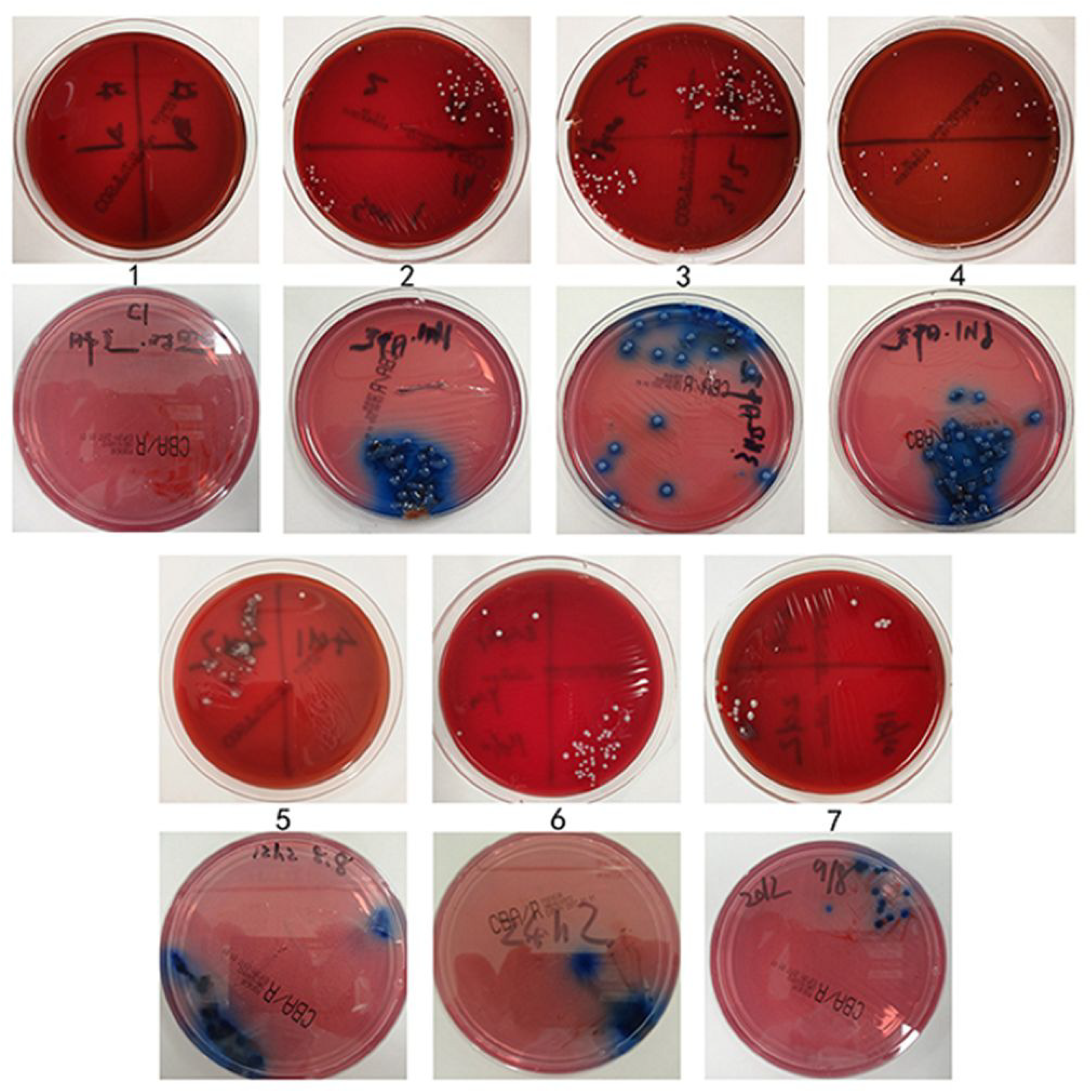
The changes of the mouse body weight. The values were presented as means ± standard deviation (S.D.) for 10 per group. *, *P* <0.05 (There were significant differences compared with the control group).

### 3.4 PCR results of isolated bacteria

The results showed that the suspected bacteria in blood culture were *S.aureus* and *K.pneumoniae.* The electrophoresis results were shown in Figure 2. The mice were confirmed to be infected by their activities, weight changes and PCR results.

**Fig 2.**
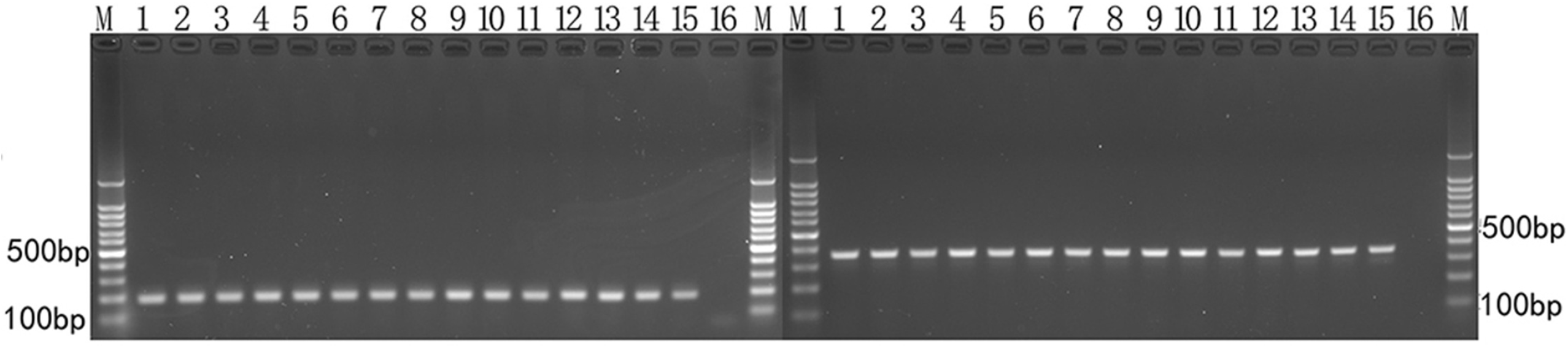
A. The results of the whole blood culture at different time after injection in mice. 1 represented the blood culture result of the control group. 2-7 represented the results of 0.5 h, 3 h, 6 h, 12 h, 24 h and 48 h after injection, respectively. Take 1 as an example, the above was the cultivation result of the blood agar, the following was the cultivation result of the China blue plate. B. The electrophoretic results after PCR. M represented the marker. 1-14 represented the results of 0.5 h, 1 h, 3 h, 6 h, 12 h, 24 h and 48h after infection, respectively. 15 and 16 represented the positive control and negative control, respectively.

### 3.5 Kinetic changes of nine cytokines in *S.aureus* and *K.pneumoniae* mouse infection models

The levels of 9 cytokines detected in blood collected from the three groups mice between 0 to 48 hours after infection are shown in Figure 3.

**Fig. 3.**
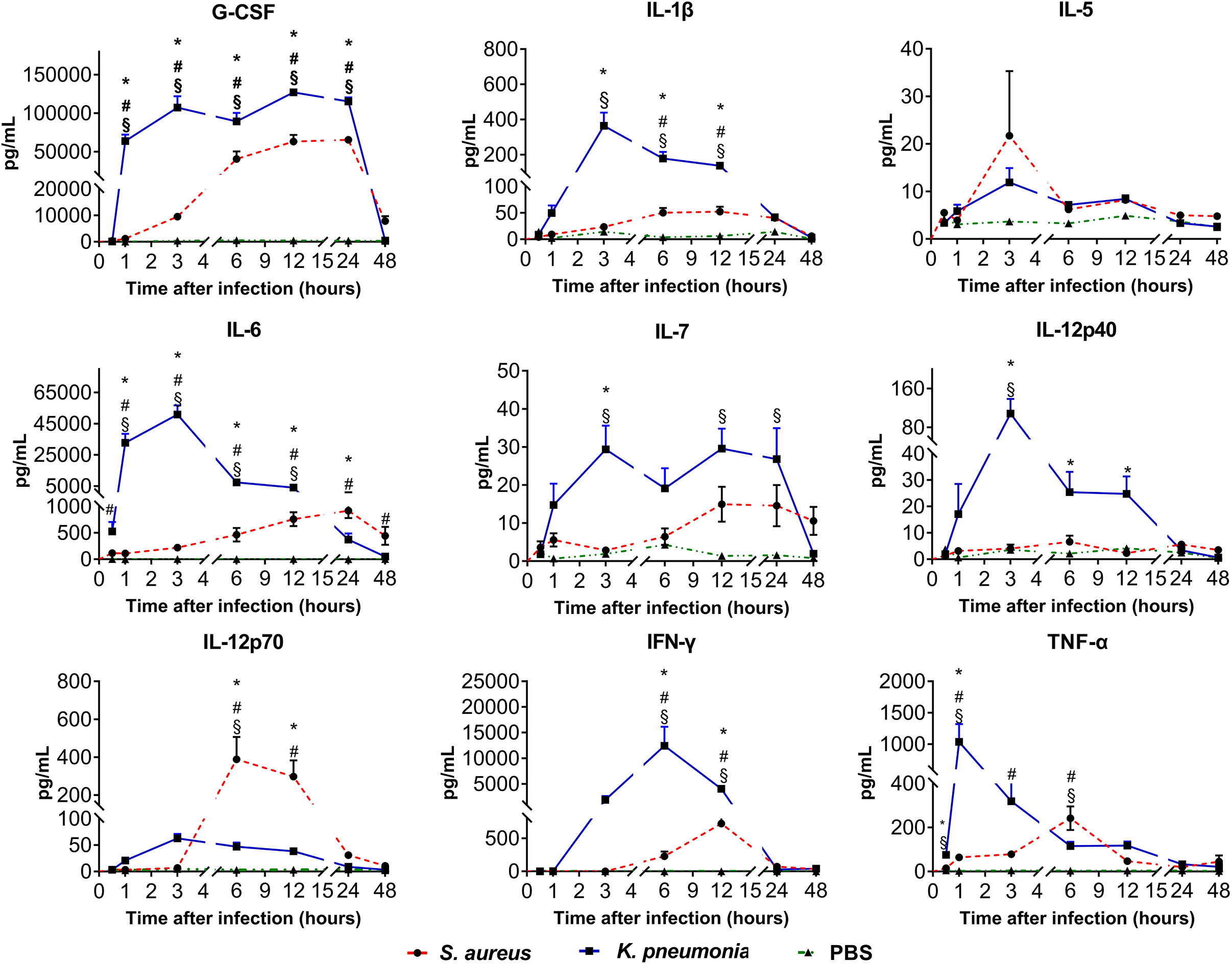
The kinetic changes of 9 cytokine in the sera of infected mouse models of S. aureus and K. pneumoniae and PBS control groups.

The 9 cytokines had started to increase at 0.5 hours after infection and decreased back to their initial level at 48 hours after reaching the highest peaks. The rate and quantity of the cytokines increased in group *K.pneumoniae* were higher than that of group *S.aureus* after infection except for IL-12p70. In the 9 cytokines, the substantial change of G-CSF is the same as IL-6 between 0.5 and 6 hours, and after the latter time point, the level of IL-6 dropped rapidly whereas that of G-CSF remains high. In contrast, there were no significant changes in the levels of the cytokines in control group mice during the experiment.

### 3.6 Discrimination of 6 validated cytokines in human serum samples among the infected *with S. aureus* and *K. pneumoniae*

We defined 0 to 6 hours after infection in mice as the early stage of the bacterial infection. In this stage, the differences were found for G-CSF between any two groups in the three groups at 0.5h, 1h, 3h, 6h, and for IL-6, they were at 1h, 3h, and 6h. For TNF-α, the difference was at 1h, for IL-12p70, IFN-γ and IL-1β, they were at 6h, (*P*<0.05) whereas for IL-5, IL-7and IL-12p40 (*P*>0.05), differences were not found at any time point. Then the 6 cytokines showed significant differences in between the three groups were chosen to be validated using clinical samples. The results obtained by software analyzed the mean and standard deviation of the graph, and calculated the differences between the three experimental groups, as shown in Figure 4.

**Fig. 4.**
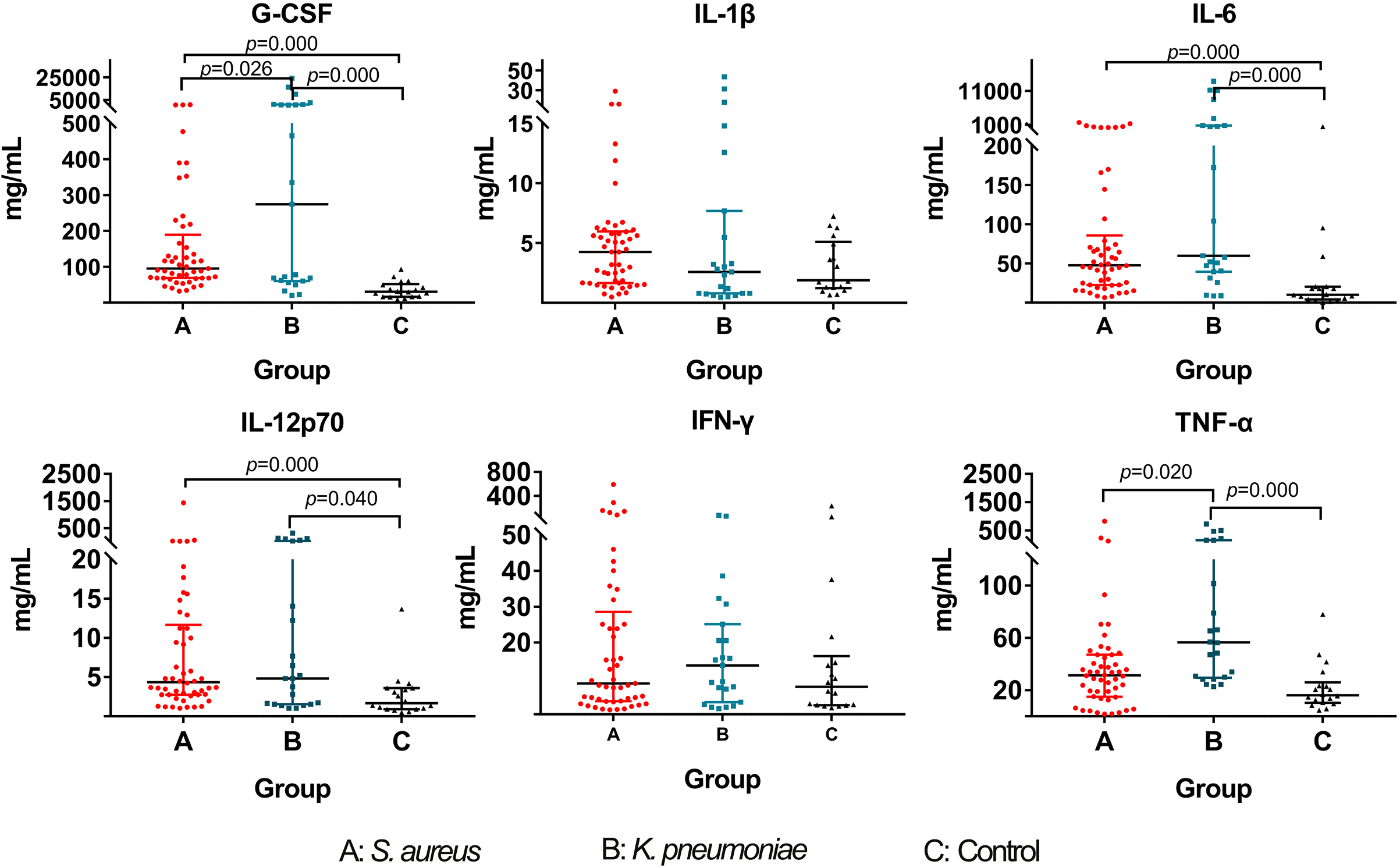
Differences of 6 validated cytokines in human serum samples among the infected with *S. aureus* and *K. pneumoniae* and PBS

### 3.7 The selected cut-off values for the potential biomarkers for bacterial bloodstream infection

The ROC curves of IL-1β, IL-6, IL-12p70, G-CSF, IFN-γ and TNF-α for diagnosing bacterial bloodstream infection are shown in Figure 5a. The AUCs calculated from the ROC curves were 0.9051 (95% Confidence interval (CI) = 0.8375–0.9767; *P* <0.0001, bootstrap=1000, the same below) for G-CSF, 0.8227 (95% CI = 0.7026-0.9427; *P* <0.0001) for IL-6, 0. 7309 (95% CI = 0.5959-0.8659; *P*=0.0064) for IL-12p70, 0.6875 (95% CI = 0.5478-0.8272; *P*<0.0270) for TNF-α, and the AUC of IL-1β and IFN-γ was 0.5875 (95% Confidence interval (CI) =0.4500-0.7250, *P* =0.2519) and 0.5069 (95% CI =0.3339-0.6800, *P* =0.9393) respectively.

**Fig. 5.**
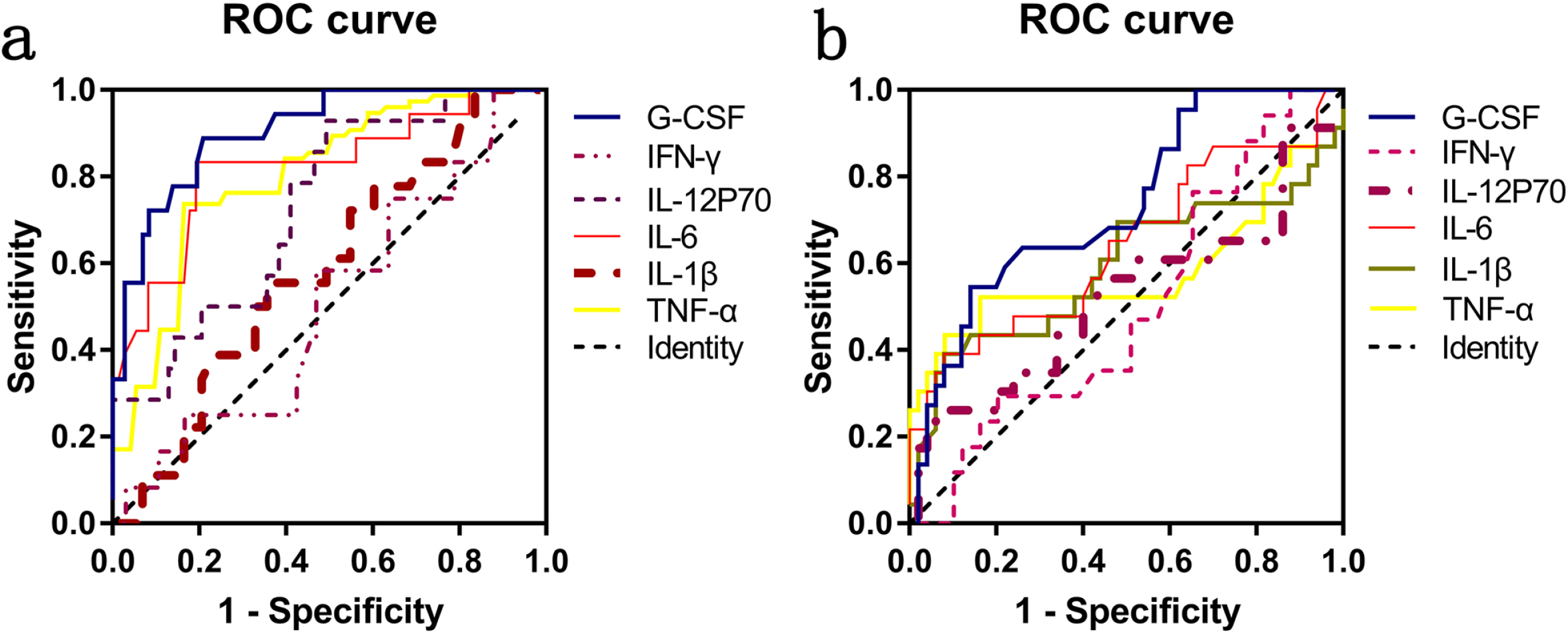
ROC curve for 6 cytokines for distinguishing bacterial infection from PBS (a) and *S. aureus* from *K.pneumoniae* infection (b).

The selected cut-off values for G-CSF, IL-6 etc. and diagnostic accuracy for bacterial bloodstream infection are shown in Table 1.

**Table 1.**
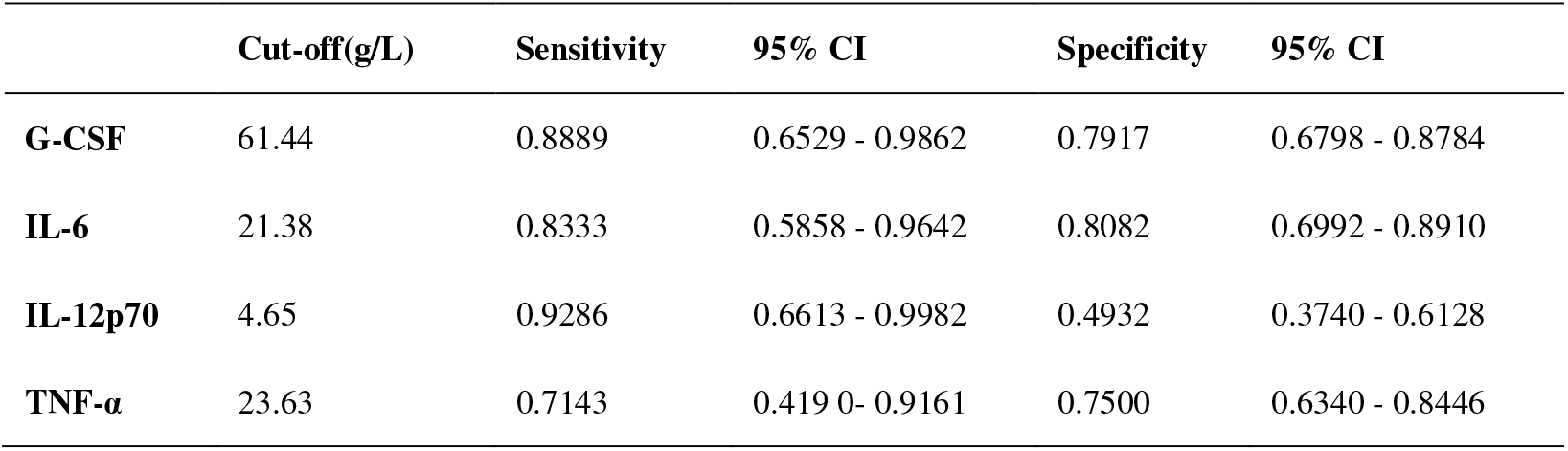
The selected cut-off values for the potential biomarkers for bacterial bloodstream infection

The ROC curves of IL-1β, IL-6, IL-12p70, G-CSF, IFN-γ and TNF-α for distinguishing *S. aureus* or *K.pneumoniae* infection are shown in Figure 5b. The AUCs calculated from the ROC curves were 0.7350 (95% Confidence interval (CI) = 0.6118-0.8582; *P* <0.05) for G-CSF and 0.6431 (95% CI = 0.4938-0.7888; *P* =0.0536) for IL-6, whereas all other biomarkers had failed to differentiate the two bacterial infections, data are shown in table 2.

**Table 2.** The P values for the potential biomarkers for distinguishing *S. aureus* or *K.pneumoniae* infection

|  | AUC | 95% CI | P value |
| --- | --- | --- | --- |
| **G-CSF** | 0.7350 | 0.6118-0.8582 | 0.0259 |
| **IL-6** | 0.6413 | 0.4938-0.7888 | 0.0536 |
| **IL-12p70** | 0.5248 | 0.3673-0.6823 | 0.7350 |
| **TNF-α** | 0.5816 | 0.4110-0.7523 | 0.2665 |
| **IL-1β** | 0.5922 | 0.4308-0.7535 | 0.2081 |
| **IFN-γ** | 0.5006 | 0.3466-0.6546 | 0.9942 |

## 4. Discussion

In recent years, the morbidity of septicemia caused by bacteria infection has increased [8]. Lack of timely diagnosis and control of initial infections, together with drug-resistance due to the overuse of antibiotics, destroys the normal flora of patients, and persistent flora disorder in the body can lead to fatality[1, 2]. Therefore, early diagnosis of bacterial infections in patients is of great significance. Specific and sensitive early biomarker for diagnosis of bacterial infection remains a challenge in current medicine, though many novel biomarker was constantly being reported.

We wonder to understand the dynamics of these cytokines associated with bacterial infection in the body, but we were unable to obtain these data in humans, so we chose a mouse model of bloodstream infection to understand this.

In the early stage of bacterial infection, the organism mainly produces innate immune response. The surface material of bacteria can be combined with the PRR of immune cells, and the transcription and expression of inflammatory factors can be activated through cell signal transduction pathways such as MyD88[9]. Due to the difference in the cell wall composition of gram-negative and positive bacteria, the innate immune response caused by this is different, and the inflammatory factors is different. Therefore in this experiment we also aimed to distinguish the two types of bacterial infection, so that clinicians can make more informed treatment decisions.

Zarkesh, M, etc. thought IL-6, CRP, WBC, and absolute neutrophil count have diagnostic value to predict serious bacterial infection [10]. Du WX etc. thought the Interleukin 35 was a novel candidate biomarker to diagnose early onset sepsis [11]. Presepsin (sCD14-ST) and others [12, 13] was also assessed as a biomarker of infection. Of all the new infectious biomarkers, only IL-6 is thought to be more helpful for the diagnosis. It is possible to need a panel biomarker to decide the bacterial infection [14, 15]. Interleukin 6 (IL-6) is an interleukin that acts as both a pro-inflammatory cytokine and an anti-inflammatory myokine[16], and it is secreted by T cells and macrophages to stimulate immune response, plays a role in fighting infection, so it is often used clinically as an inflammation biomarker.

In the mice model, the performance of IL-6 is consistent with the literature [10], it could discriminate infection and no-infection. Interestingly, our results showed that the variation tendency of G-CSF is not only very similar to IL-6, but it also rose more strongly than IL-6 in mouse serum. During post-infection analysis, the G-CSF reached its peak rapidly at 1 h after infection and lasted 24h, then it was rapidly descended to the initial level until 48h. The body response of the mice to *S. aureus* infection was weaker than that of *K. pneumoniae*, and there are significant differences between the three experiment groups (*P*<0.01). G-CSF is a glycoprotein that stimulates the bone marrow to produce granulocytes and stem cells and release them into the bloodstream[17]. It can be produced by endothelium, macrophages, and a number of other immune cells from different tissues. Functionally, it could not only act as a cytokine but also a hormone, and it stimulates the survival, proliferation, differentiation, and function of neutrophil precursors and mature neutrophils. Based on its performance in the study, we believe that G-CSF is a better infectious biomarker than IL-6.

Changes in the four cytokines IL-1β, IL-5, IL-7 and IL-12p40 within 6 h after infection in the animal model were not significantly different between the two bacteria, and mice appeared to have weaker responsive to *S.aureus* than *K. pneumoniae*,. This results had first led us to suspect that if the variance was caused by different number of bacteria injected between *S.aureus* and *K. pneumoniae*. However, the dynamics curve of IL-12p70 dispelled the conjecture. IL-12p70 level of the mice after infection with the same *S.aureus* dose was significantly higher than that of *K. pneumoniae*, so we then speculated that this might be related to the bacterial surface structure. IL-12p40 and IL-12p70 belong to different subunits of IL-12. The performance of these two cytokines in this experiment show there exists influence each other in the opposite manner [18, 19].

The trend of IFN-γ and TNF-α in this animal model shows that they respond more slowly to bacterial infections than other cytokines.

48 hours after mice were infected with bacteria, the dynamic changes of inflammatory cytokines (such as G-CSF) were remarkably different between infectious groups and non-infectious group. There were also differences between the group of gram-positive bacteria and the group of gram-negative bacteria. However, other cytokines did not show the same difference.

When clinical control samples were collected, some of which with CRP> 6.0 mg / ml or PCT> 0.8 mg / ml or IL-6> 5.9 mg / ml were excluded because the positive rate of clinical blood culture was only 10-20%, therefore, although some cultures are negative for blood cultures, some may still be infected.

In selecting theory validated cytokines, we were aiming to find biomarkers that could discriminate infected and non-infected patients, as well as gram-negative and positive bacteria, in the early stage of infection. So, we selected 6 cytokines that differed among the 3 groups of mice within 6 hours of post-infection in the animal model.

The clinical validation results showed that IL-12p70, G-CSF and IL-6 had significant differences in the infection and non-infection population. Although TNF-α was a factor that can distinguish *S.aureus* group from *K. pneumoniae* group. *S.aureus* group and SIRS group cannot be distinguished from each other. Only G-CSF had the power to differentiate infection/SIRS group and also *S.aureus* group and *K. pneumonia* group.

From the ROC curves that distinguish between infected and no infected, we can see. The AUC of G-CSF was the highest among the six cytokines (AUC = 0.9051), which was superior to the IL-6 (AUC = 0.8227). In *S.aureus* group and *K. pneumoniae* group ROC curves. AUC of G-CSF = 0.7350 *(P* <0.05). This means that unlike other cytokines, G-CSF was the only one showed significant differences in all testing groups.

The change of these nine cytokines in the animal model reflects the body’s kinetic changes after bacterial infection. In the mouse model, with the progression of infection time, the level of cytokines is also undergoing major changes.

In contract, the clinical samples from patients are taken at a single, specific time point, and we do not know exactly when the patient is infected. Therefore, it was not possible to accurately compare the results obtained from the mice model and the patients. Nevertheless, our statistical analysis can still reflect the changes of inflammatory factors in the patients after infection to a certain extent.

### Conclusions

Based on the kinetic changes of these nine cytokines in animal models and the clinical validation of six of the cytokines, G-CSF is found to be suitable as a potential biomarker for early detection of infection and as a possible discriminator between Gram-negative and positive infection. We will verify the changes of these factors in other common clinical gram-positive and negative bacteria in future trials. More clinical samples are warranting for future validation.

This study is the first part of this series of studies. We decide to observe inflammatory cytokines with other types of bacteria in the future study.

## Declarations

### Ethics approval and consent to participate

This study was approved by the Chinese PLA medical ethics committee and the experimental animal ethics committee.

### Consent to publish

All authors have agreed to publish this manuscript.

### Availability of data and materials

The data and materials included in this manuscript are available.

### Authors’ Contributions

Chengbin Wang and Jiyong Yang designed and supervised the experiment; Shang He contributed to preparation of bacteria and manuscript; Ming Yang contributed to animal experiment and measurement of cytokines; Xinjun Li and Yating Ma contributed to collection of clinical samples and patients’ clinical informtaiton; Chen Chen contributed to revision of manuscript; Chi Wang contributed to data analysis.

## Acknowledgements

We would like to acknowledge the guidance from microbiological specialist Professor Yanping Luo from our department.

